# Advancing Therapeutic Solutions: Poloxamer-based Thermosensitive Injectable Hydrogels containing a Self-assembling Peptide for *In situ* Gelation in an Osteoarthritis Murine Model

**DOI:** 10.1101/2025.04.30.651282

**Authors:** S.Sana Sayedipour, Timo Schomann, Sanne M. van de Looij, Somayeh Rezaie, Yolande F.M. Ramos, Tina Vermonden, Louise van der Weerd, Ingrid Meulenbelt, Luis J. Cruz

## Abstract

This study presents the development and characterization of a novel thermosensitive injectable hydrogel designed to enhance the biomechanical properties of poloxamer 407 (P407) through the incorporation of a self-assembling peptide. The primary objective was to engineer a formulation that rapidly gels following intra-articular (i.a.) injection, exhibits improved mechanical strength, and enables sustained release of embedded therapeutic cargo.

Gelation time assays demonstrated that the P407-peptide formulation solidified more quickly than P407 alone at equivalent concentrations. Rheological analysis revealed a 1.5 kPa increase in storage modulus in the hybrid hydrogel, confirming improved mechanical integrity. *In vitro* biocompatibility was assessed using human chondrocytes, with MTS assays and LIVE/DEAD staining indicating no cytotoxicity across tested concentrations.

To evaluate *in vivo* applicability, a near-infrared fluorescent (NIRF) dye was incorporated into the hydrogel and injected intra-articularly into an osteoarthritis (OA) mouse model. The labeled formulation allowed for successful tracking and demonstrated localized gelation, supporting its suitability for site-specific, sustained delivery.

Overall, the P407-peptide hydrogel offers a promising platform for i.a. therapeutic applications, combining injectability, rapid thermoresponsive gelation, mechanical reinforcement, and controlled release behavior, making it well-suited for regenerative medicine and OA treatment.

## 1. Introduction

Injectable hydrogels have gained significant attention as *in situ* drug delivery systems due to their tunable physical properties, minimally invasive administration, and ability to provide sustained release of therapeutics at the target site [1-4]. Such properties make thermosensitive hydrogels particularly suitable for intra-articular (i.a.) administration in osteoarthritis (OA) treatment, where encapsulation of therapeutic cells or drugs within the hydrogel matrix enables targeted delivery directly into the joint space [5]. In particular, thermosensitive hydrogels are attractive drug delivery systems, remaining liquid at lower temperatures during injection and rapidly forming a gel at physiological temperatures. This temperature-triggered transition enables localized, sustained release of therapeutics, reducing systemic toxicity and minimizing the frequency of administration[6, 7]. Among them, Poloxamer 407 (P407) has been widely studied as a result of its unique thermo-reversible gelation properties, low toxicity and sustained drug release characteristics. P407 is a triblock copolymer with a center block of hydrophobic polypropylene oxide (PPO) between two hydrophilic polyethyleneoxide (PEO) lateral chains [8-10]. Upon dissolution in water, P407 molecules self-assemble into micelles with a hydrophobic polypropylene oxide (PPO) core and a hydrophilic polyethylene oxide (PEO) shell. When the polymer is sufficiently concentrated and the temperature rises beyond the critical gelation temperature (CGT), these micelles organize into a firm network, leading to the formation of a semi-solid gel depot [11]. Despite the multiple benefits of P407, proper application of this gel faces difficulties due to its rapid dissolution in aqueous media and low mechanical strength [12]. To overcome these limitations, various strategies have been explored, including the incorporation of pharmaceutically active ingredients, salts, excipients, and functional biomolecules into P407 formulations, which can modulate the sol–gel transition temperature (T_sol–gel_) and enhance hydrogel performance[13, 14]. Among these approaches, the incorporation of self-assembling peptides has emerged as a particularly effective strategy to improve the structural integrity and functional properties of P407-based hydrogels[15]. These peptides can physically interpenetrate or chemically interact with the hydrogel network, improving cargo affinity, inhibiting burst release, and prolonging drug release, thereby making the system more suitable for controlled delivery applications [16].

In this study, a thermosensitive hydrogel based on P407 containing a amphiphilic self-assembling peptide was developed. The effect of the polymer concentration on the gelation behavior in combination with the peptide was analyzed. *In vitro* studies on the human chondrocyte cell line C28/I2 were performed to assess the toxicity of a new formulation of P407. Additionally, We performed *in vivo* experiments using a mouse model of OA to evaluate the thermosensitive injectability, localization, and sustained release capabilities of a peptide-modified P407 hydrogel. For this purpose, we incorporated the near-infrared fluorescent (NIRF) dye IR780 into the hydrogel and administered it via i.a. injection into the knee joint d OA mice model. Our findings support the hydrogel’s potential as an injectable delivery system for clinical applications in OA therapy.

## 2. Materials and methods

### 2.1. Materials and reagents

Commercial P407 (MW= 12,600Da) and phosphate-buffered saline (PBS) were produced by Fresenius Kabi GmbH (Graz, Austria). Dulbecco’s phosphate-buffered saline (DPBS), Dulbecco’s modified eagles’ medium (DMEM, high glucose, with Glutamax™), fetal bovine serum (FBS), penicillin, and streptomycin were purchased from Life Technologies (Breda, the Netherlands). 3-(4,5-Dimethylthiazol-2-yl)-5-(3-carboxymethoxyphenyl)-2-(4-sulfophe-nyl)-2H-Gels 2022, 8, 44 11 of 16 tetrazolium (MTS, Promega, Madison, WI, USA) and a calcein-AM/ethidium homodimer-1 LIVE/DEAD® assay kit (Invitrogen) were obtained from Carlsbad, CA, USA

### 2.2. Self-assembling Peptide

Self-assembling peptide with sequence Palmitoyl-WKGNNQQNYQQ, designed by Dr. Luis J. Cruz (Department of Radiology, Leiden University Medical Center, Leiden, the Netherlands) and was synthesized using solid-phase peptide synthesis (SPPS) employing the Fmoc strategy[17] and provided by GL Biochem Ltd. (Minhang, Shanghai, China).

### 2.3. P407-peptide hydrogel preparation and characterization

#### 2.3.1. P407-peptide hydrogel preparation

P407-peptide composite hydrogel was prepared using the ‘cold’ method[18]. To prepare 30% hydrogel(w/v), 3gr of poloxamer powder and 100mg of peptide were gently dissolved in 10mL of sterilized ultrapure water. The mixture was then stirred at 4°C overnight to ensure that all the powder dissolved completely, resulting in a clear solution.

#### 2.3.2. Time-to-gelation assay

The time-to-gelation temperature of P407 and P407-peptide hydrogels was evaluated using the tube inversion method. The peptide was incorporated into the P407 solution at a concentration of 1% (w/v). For each formulation, 1 mL of hydrogel was transferred into a test tube and incubated in a water bath at either room temperature (RT, 25 °C) or 37 °C. The sol–gel transition temperature was defined as the point at which the solution no longer flowed upon tube inversion, indicating the formation of a gel. Each measurement was performed in triplicate.

#### 2.3.3. Rheological characterisation

The rheological properties of 25% P407 and 25% P407 containing 1% (w/v) self-assembling peptide were evaluated using a Discovery HR-2 Rheometer (TA Instruments, Etten-Leur, The Netherlands) equipped with a Peltier plate for precise temperature control and a solvent trap to minimize sample evaporation. Both samples were measured using a 20 mm diameter aluminium plate-plate geometry and an initial gap of 300µm. Hydrogel formulations were prepared, and 100 µL sample were deposited under the geometry for measurements. Data were processed with TRIOS Software version 5.1. The appropriate strain for all samples was determined by formation of the hydrogel formulations at 37°C for 30 min, followed by an oscillation amplitude with increasing strain from 0.01% to 100% at a frequency of 1 Hz. Gelation kinetics were then assessed by measuring the storage (G’) and loss (G’’) moduli during a temperature ramp ranging from 4°C to 37°C at a rate of 1.0°C.min^-1^, using a frequency of 1 Hz and a strain of 0.1%. Next, the samples were kept at 37°C for 15 min, before cooling down to 4°C at a rate of 1.0°C.min^-1^.

#### 2.3.4. Morphology of P407, self-assembled peptide network and P407-peptide hydrogels by scanning electron microscopy (SEM)

A 25% (w/v) solution of P407 and an equivalent concentration of 25% P407 containing 1% (w/v) self-assembling peptide were frozen at −80 °C and subsequently lyophilized to examine hydrogel morphology. After freeze-drying, small pieces of each sample were cut, mounted on aluminum stubs, and sputter-coated with a platinum layer using a Cressington 208HR sputter coater (Cressington, Watford, UK). The surface structure and pore size were then analyzed using a NanoSEM 450 scanning electron microscope (FEI, Tokyo, Japan).

#### 2.3.5. Dynamic light scattering (DLS) measurements

Samples with different concentrations of P407 and P407-peptide (1, 2, 3, 4, 5, 7.5 and 10%) were prepared in MilliQ H_2_O (Merck-Millipore, Burlington, MA, USA) to study the properties of P407 micelles. Approximately 800mL of the samples were then transferred to cuvettes for measurement using a Zetasizer (Nano ZS, Malvern Ltd., Malvern, UK).

#### 2.3.6. P407 and P407-peptide hydrogel degradation

The release kinetics of P407-peptide hydrogel were assessed using the direct release method, with NIRF dye IR780 as the loaded cargo[12, 19]. In brief, 50ng of IR780 was mixed with 1mL of 25.0% (w/v) P407 and P407-peptide hydrogels in 15mL tubes at 4°C. The tubes were then incubated at 37°C for 5min to allow the formation of a stable gel. After gel formation, 4mL of pre-warmed PBS was gently added on top of the gel layer, and the tubes were placed in a shaking water bath at 37°C and 30rpm. At specific time points (1, 2, 4, 8, 12, 24, 48, 72, 96, 120, 144, and 168 hr), 1mL of supernatant was collected from each tube, and the volume was replaced with fresh PBS pre-warmed to 37°C. The released IR780 was then quantified using SpectraMax® iD3 Multi-Mode Microplate Readers (Marshall Scientific, Hampton, NH, USA).

Additionally, the degradation of the hydrogels was evaluated. Briefly, 1mL of hydrogel was placed in 15mL falcon tubes and incubated at 37°C for gel formation. The initial weight of the tube with hydrogel was recorded as 100% of the gel weight. Following this, 4mL of pre-warmed PBS was added on top of the hydrogel. At predetermined time points corresponding to those in the release measurement, the entire volume was removed, and the weight was measured to calculate the amount of hydrogel degradation.

### 2.4. *In vitro* experiment

#### 2.4.1. Cell culture

The human chondrocyte cell line C28/I2 was cultured in 75cm2 flasks containing a 1:1 mixture of DMEM and F12 medium with 10% (v/v) FBS at 37°C and 5% CO_2_. The medium was refreshed every 48hr. Once the cells reached 80-90% confluence, they were subcultured for experiments.

#### 2.4.2. Cell metabolic assay (MTS)

The effect of P407 and peptide on C28/I2 cells was evaluated using the MTS colorimetric method[20]. Cells were seeded in a 96-well plate at a density of 5×10^3^ cells per well and incubated at 37°C for 24hr. Subsequently, the cells were treated with various concentrations of P407-peptide hydrogel (w/v) for 24, 48, and 72 hr. Untreated cells served as negative control, while cells treated with 50 % DMSO served as positive, cytotoxicity control. After each time point, the old medium was carefully removed and replaced with 1mL of fresh medium plus 20μL of [3-(4,5-dimethylthiazol-2-yl)-5-(3-carboxymethoxyphenyl)-2-(4-sulfophenyl)-2H-tetrazolium (MTS) in each well. After 1h of incubation at 37**°**C, the optical density was measured using a spectrophotometer at λ_ex_ 590nm.

#### 2.4.3. Cell viability (LIVE/DEAD® assay)

The effect of 25% P407-peptide hydrogels on the viability of C28/I2 cells was evaluated using a LIVE/DEAD® assay at 1, 3, and 7 days after culture. A total of 2×10^4^ C28/I2 cells/well were seeded on top of the P407-peptide hydrogels in a 48-well plate and incubated for 1, 3 and 7 days. After removing the supernatant, an aliquot of the assay solution containing 4µM EthD-1 (ethidium homodimer-1) and 2 M calcein AM was added to the cells. Following a 25min incubation at RT, the samples were observed using an Andor Dragonfly 200 spinning disc confocal microscope (Andor Technology, Belfast, UK) with excitation filters of 450–49nm (green, Calcein AM) and 510–560 nm (red, EthE-1). Living cells were visualized as green and dead cells were visualized as red.

### 2.5. *In vivo* experiment

#### 2.5.1. Animal model

To evaluate the concept of our thermosensitive, injectable P407-peptide hydrogel and its degradation and steady release of the sample dye, we performed animal experiments using an OA mouse model based on the destabilization of the medial meniscus (DMM). For the experiment, a total of eighteen 12 weeks-old male C57BL/6J mice were purchased from Charles River Laboratories (Charles River, Chatillon-sur-Chalaronne, France). The animal procedures were all conducted at the Leiden University Medical Center and were approved by the Animal Welfare Committee (IvD) under number AVD1160020171405 -PE.18.101.005. All mice were housed in groups in polypropylene cages on a 12-hour light/dark cycle with unrestricted access to standard mouse food and water. Surgical destabilization of the medial meniscus was performed on the right knee joint as described before[21]. Three weeks after the surgery, DMM mice were randomly divided into 3 groups, receiving 6µL of either saline plus 50ng of IR780, P407 loaded with IR780, or P407-peptide loaded via i.a. injection. These mice were sacrificed 34 days after the injections, and their limbs were and studied by means of fluorescence imaging for signs of scaffold degradation.

#### 2.5.2. *In vivo* imaging

To evaluate the degradation and stability of the thermosensitive hydrogel mixed with IR780, NIRF imaging was assessed using the Pearl Impulse Imaging System (Li-Cor, Lincoln, NE, USA). Mice were anesthetized using isoflurane (4–5% induction, 1–2% maintenance) and imaged at 1, 7, 14, 21, 28, and 34 days post i.a. injection. Imaging was performed in the near-infrared (NIR) 800nm fluorescence channel, corresponding to the emission wavelength of IR780. A fixed exposure time and gain setting were used across all time points to ensure consistency in image acquisition. Fluorescence intensity was quantified using Image Studio Software (Li-Cor Biosciences). A region of interest (ROI) was manually defined around the knee joint to measure the total radiant efficiency. To correct for background fluorescence, an ROI was selected in a non-fluorescent region, and the signal was normalized accordingly.

### 2.6. Statistical analysis

For statistical analysis, GraphPad Prism 8.1.1 software (GraphPad Software, San Diego, CA, USA) was utilized. All data are expressed as the mean ± standard deviation (SD) of 3-5 independent repeated experiments, unless otherwise stated. Statistical significance was determined using a Student’s *t*-test, unpaired, Mann-Whitney U test, and two-way analysis of variance (ANOVA). In all analyses, a p-value ≤ a is considered an indicator of statistical significance and is expressed as: * p ≤ 0.05, ** p ≤ 0.01, *** p ≤ 0.001, **** p ≤ 0.0001.

## 3. Results

### 3.1. Characterization and gelation time of P407-peptide hydrogel

P407 hydrogels exist in a fluid state below their critical gelation temperature (CGT), allowing them to flow and be easily injected. Upon increasing the temperature, these hydrogels undergo a sol–gel transition and form a semi-solid hydrogel (**Figure 1**). To assess the influence of polymer concentration on gelation behavior, hydrogels containing varying concentrations of P407 (10, 15, 20, 25, and 30%) were prepared. Gelation times were recorded at room temperature (RT, 20 °C) and physiological temperature (37 °C)(**Table 1**). At concentrations of 10% and 15%, P407 hydrogels remained liquid at both RT and 37 °C, indicating that a minimum threshold of ≥18% is required for thermosensitive gelation, consistent with previous reports [22]. At 25% concentration, the P407 hydrogel exhibited gelation times of 305±5s at RT and 44±1.7s at 37 °C. Increasing the P407 concentration further decreased gelation times, showing rapid gelation even at RT, which posed practical challenges such as needle clogging during injection.

**Table 1.** Gelation time of P407 and P407-1% peptide hydrogels at various polymer concentrations.

| Formulation | Time to Gelation at RT (20°C) | Time to Gelation at 37°C |
| --- | --- | --- |
| 10% P407 | - | - |
| 15% P407 | - | - |
| 20% P407 | 467.6±15.7 | 89.3±4 |
| 25% P407 | 305±5 | 44±1.7 |
| 30% P407 | 125.3±7.6 | 24±1.7 |
| 10% P407 + 1%Peptide | - | - |
| 15% P407+ 1%Peptide | - | 98.3±2.8 |
| 20% P407+ 1%Peptide | 365.3±4.1 | 68±3.6 |
| 25% P407 + 1%Peptide | 180.3±8.5 | 32±2* |
| 30% P407+ 1%Peptide | 110.6±12.5 | 18±2.5 |

Reported in seconds measured at room temperature (RT) and at 37°C. Results are shown as means of a triplicate measurement ± standard deviation (SD). \* Chosen for subsequent experiment.

**Figure 1:**
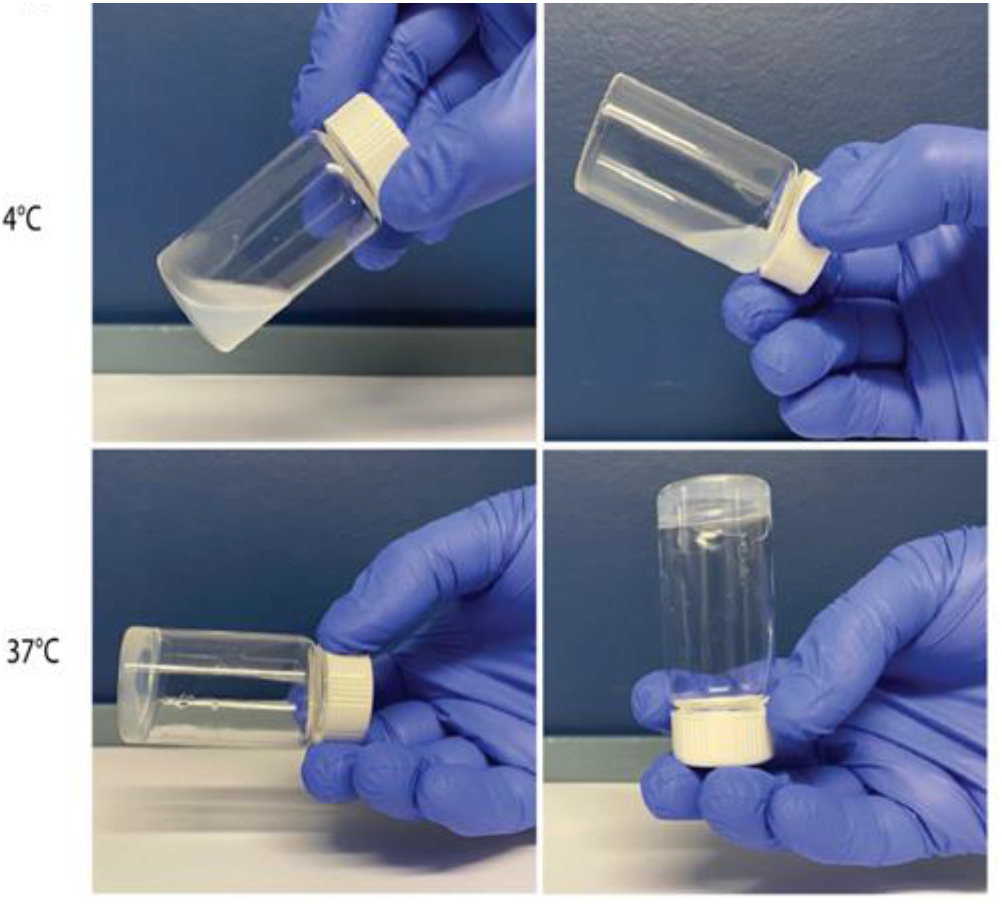
P407 hydrogels. The optical images of 25% P407 hydrogel sol–gel transition at 4°C and 37°C.

To enhance the structural stability of the P407 hydrogel while maintaining injectability, we incorporated varying concentrations (1, 2, 3, 4, and 5%) of a self-assembling peptide (Palmitoyl-WKGNNQQNYQQ) into P407 formulation. Peptide concentrations greater than 1% led to the formation of bulk solid aggregates that clogged the injection needle and prevented thermosensitive gelation. Therefore, 1% peptide was identified as the optimal concentration for incorporation.

We next compared the gelation behavior of the optimized P407 + 1% peptide hydrogel relative to P407 alone by performing time-to-gelation assays (**Table 1**). Incorporation of 1% peptide into the 25% P407 formulation significantly decreased the gelation time at both 20 °C and 37 °C. Specifically, gelation times were reduced to 180.3±8.5 s at RT and 32±2 s at 37°C, compared to longer gelation times observed for the unmodified 25% P407 hydrogel. Importantly, the 25% P407 + 1% peptide formulation maintained favorable injectability, without needle clogging during administration. This contrasts with the 30% P407 hydrogel, where excessive viscosity at RT caused needle clogging and impaired handling. The acceleration of the sol–gel transition, combined with good injectability at room temperature, is particularly advantageous for intra-articular applications, as it enables rapid in situ depot formation at the target site while ensuring ease of administration. Based on these findings, the 25% P407 + 1% peptide formulation was selected for all subsequent experiments. The 25% P407 hydrogel without peptide was used in parallel as a control to directly compare the effects of peptide incorporation on the hydrogel’s physicochemical and biological properties.

### 3.2. Rheology characterization of the hydrogel

Rheological studies were conducted to assess the gelation kinetics and stiffness of 25% (w/v) P407 control and the same concentration of P407-peptide hydrogel formulations. First, gels were fully formed at 37°C and then subjected to an oscillation amplitude at a constant frequency, where the G’ and G’’ were measured when the strain increased exponentially. As visualised in **supplementary Figure 1**, both hydrogels show a sharp decline in gel strength above a strain of 0.6%. An appropriate strain before the breaking point of the gel (0.1 %) was chosen for subsequent experiments. Next, the gelation kinetics upon heating and cooling of gel formulations were assessed. Herein, the gelation temperature (Tgel) is defined as the temperature at which G’ overtakes G’’. Both formulations displayed a progressive rise of the G’ from Tgel to 37°C (**Figure 2A and B**), which can be attributed to the thermosensitive nature of the poloxamer-based polymer. In particular, P407 control showed a Tgel of 23.0 °C, while the addition of the self-assembling peptide decreased the Tgel to 20.5°C, as summarised in **Table 2**. Furthermore, at 37°C, the P407-peptide hydrogel exhibited a higher G’ of 18.1 kPa compared to 16.6 kPa for the P407 control, reflecting improved stiffness. These enhancements in mechanical properties are evident from the progressive increase in G’ during the heating cycle and support the utility of the P407-peptide hydrogel as a robust and thermosensitive platform for sustained drug delivery applications.

**Table 2:** Gelation temperature of 25% P407 control and 25%P407-peptide hydrogel formulation.

| | T at $G' > G''$ (Heating, **) | T at $G' < G''$ (Cooling, **) | Maximum $G'$<br>(kPa, *) |
| --- | --- | --- | --- |
| **25% P407** | $23.0 \pm 0.72^{\circ}\text{C}$ | $19.8 \pm 0.23^{\circ}\text{C}$ | $16.6 \pm 0.67$ |
| **25% P407-peptide** | $20.5 \pm 0.02^{\circ}\text{C}$ | $16.9 \pm 0.60^{\circ}\text{C}$ | $18.1 \pm 0.13$ |

Statistical significance between P407 and P407 with 1% peptide is shown as \* $p \leq 0.05$ or \*\* ( $p \leq 0.01$ ).

**Figure 2:**
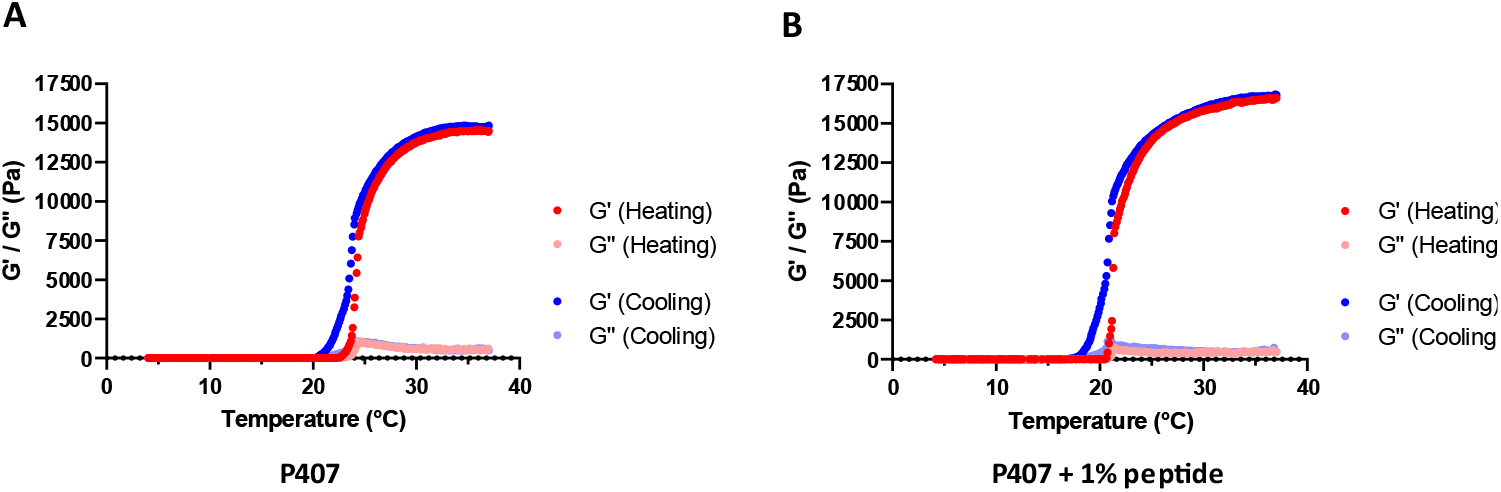
Average gelation profile of (A) P407 control and (B) P407 with 1 % self-assembling peptide gel formulations. Graphs depict the G’ and G’’ upon heating and cooling the samples between 4 and 37 °C, with each point representing the average of n = 3 measurements.

### 3.3. Morphology of 25% P407 and 25% P407-peptide

SEM revealed that 25% P407 and the corresponding P407-peptide hydrogels revealed the typical porous structure of lyophilized P407 hydrogel specimens(**Figure 3**). Pores were predominantly round and oval, with pore sizes ranging from 40±8.6μm for 25% P407 and 92±17.88μm for 25% P407-peptide as analyzed by ImageJ software. The porous morphology corresponds to the P407 micelle-based network formed during gelation, which is dependent on polymer concentration.

**Figure 3:**
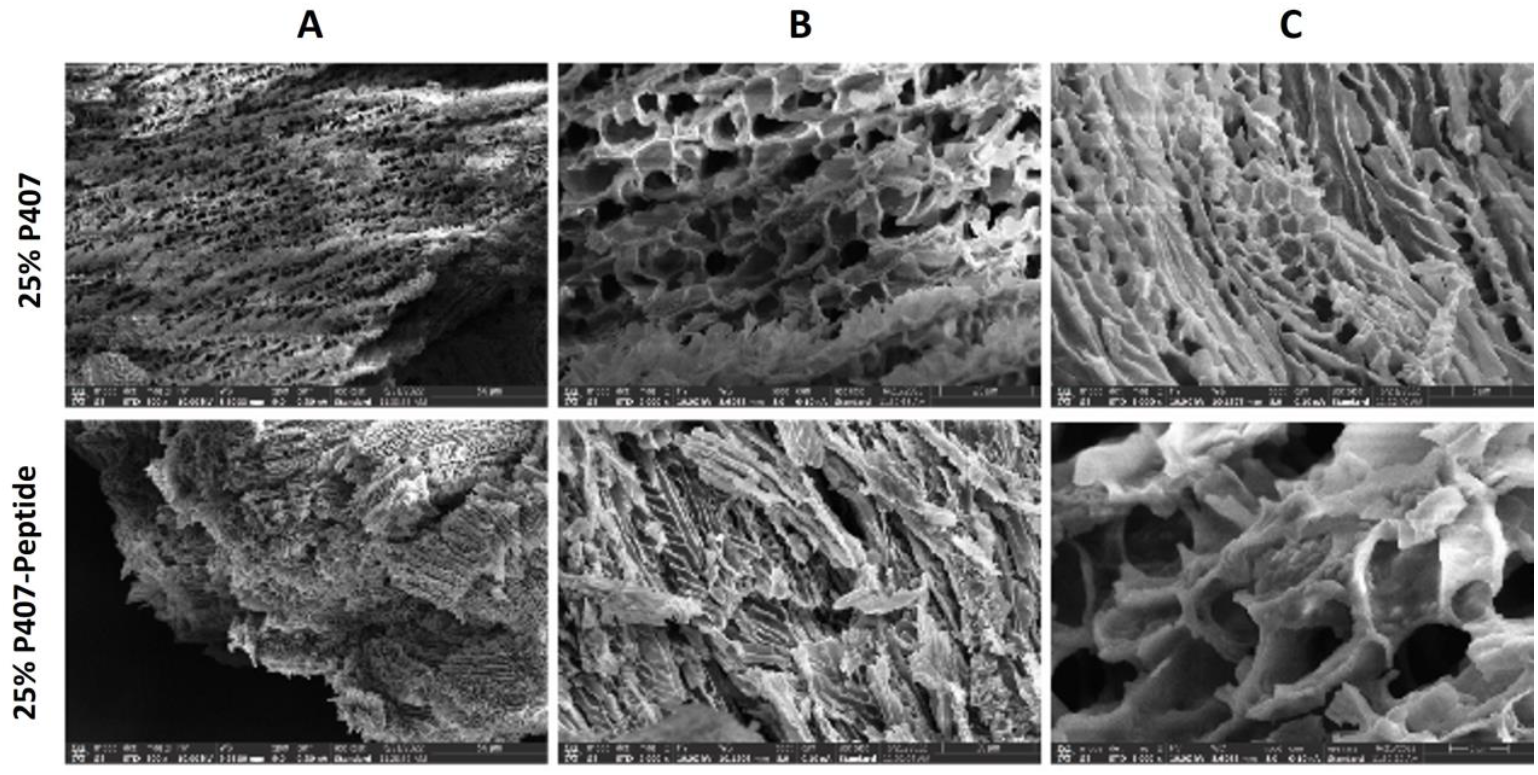
SEM images showing cross-sections of the 25% P407 and 25% P407 containing 1% self-assembling peptide hydrogels. Representative images are shown at different magnifications: (**A**) 50 nm scale bar, (**B**) 21 nm, and (**C**) 5 nm.

Dynamic light scattering (DLS) analysis (**Supplementary Figure 2**) confirmed micelle formation at 1% of P407. At this concentration, the average micelle size was 6.5 ± 0.38 nm for 1% P407 and 136.1 ± 10.59 nm for P407-peptide. Increase in polymer concentration led to a corresponding increase in average particle size in both hydrogel types. For that matter, more pronounced effect observed in the P407-peptide formulation. The larger particle size in the peptide-containing formulation correlated with the formation of a more open and expanded porous network, as observed in the SEM images.

### 3.4. 25% (w/v) P407-self-assembling peptide hydrogel enhances hydrogel stability in aqueous solutions

The stability of the 25% (w/v) P407-self-assembling peptide hydrogel was evaluated relative to same concentration of P407, by analyzing the release kinetics of IR780 and the hydrogel’s degradation over time.

The degradation profiles of the hydrogels are shown in **Figure 4A**. In the first 24hr, no statistically significant difference was observed between the two groups. At 48hr time point, a moderate but statistically significant difference in degradation became apparent (p ≤ 0.05), with the P407 hydrogel exhibiting a faster dissolution rate. Complete degradation of the P407 hydrogel occurred by day 7 (168hr), whereas the P407-peptide hydrogel remained intact until day 9(216hr).

**Figure 4:**
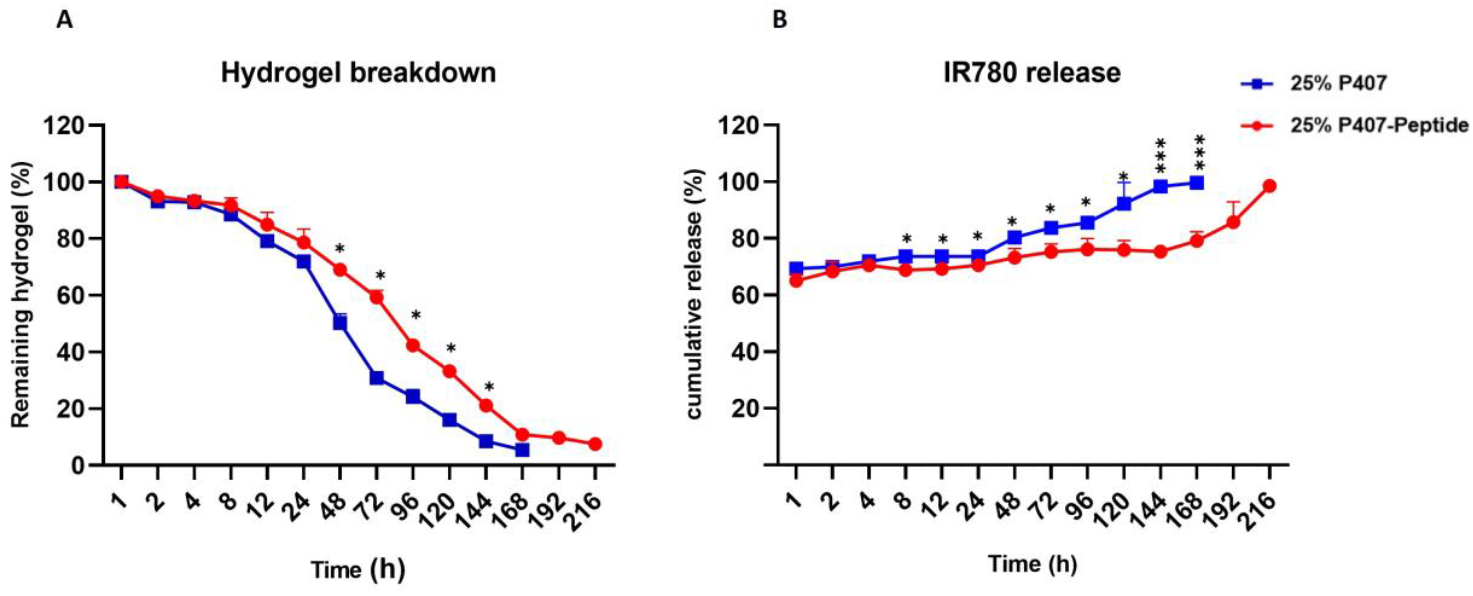
Stability and IR780 Release Profile of 25% P407 and P407-Peptide hydrogels. **(A)** Cumulative release of IR780 dye from hydrogels over time. **(B)** Hydrogel degradation over time. The data is presented as the mean ± SD, n = 3. * p ≤ 0.05, *** p ≤ 0.001, ****p ≤ 0,0001.

The cumulative release of IR780 from both hydrogels is shown in **Figure 4B**. During the first 24hr, approximately 65% of the initially loaded IR780 was released from both hydrogels. However, beyond 24hr, the release rate from the P407-peptide hydrogel progressively declined, with the remaining dye being completely released over a 9-day period. In comparison, the P407 hydrogel released the dye in just 7 days. Statistically, significant differences in IR780 release between the two hydrogels were observed at several time points. Between 72 and 120hr, moderate differences were noted (p ≤ 0.05), while highly significant differences (p ≤ 0.001) occurred at 144 and 168 hr, where the P407 hydrogel released 23.07% and 20.45% more dye than the P407-peptide hydrogel, respectively. The slower initial release rate of the NIRF dye from the P407-peptide hydrogel may be due to affinity of IR780 with the peptide, but the hydrogel dissolution had a dominant effect on IR-780 discharge.

### 3.5. *In vivo* degradation study

To assess the *in vivo* degradation behavior and retention capacity of the thermosensitive injectable hydrogels, 25% (w/v) P407 and P407–self-assembling peptide hydrogels loaded with the IR780 were injected into the right knee joints of DMM mice. PBS containing free IR780 was used as a control. Each formulation contained 50 ng of IR780.

Fluorescence intensity measurements were performed at multiple time points using the PEARL imaging system. Representative images (**Figures 5A–C**) show that free IR780 in PBS was rapidly cleared from the joint, with minimal signal detectable by day 7. In contrast, both the P407 and P407/peptide hydrogel groups maintained stronger fluorescence signals over time. The highest fluorescence retention was observed in the P407/peptide hydrogel group, followed by the P407 hydrogel group, with the PBS group displaying the weakest signal.

**Figure 5.**
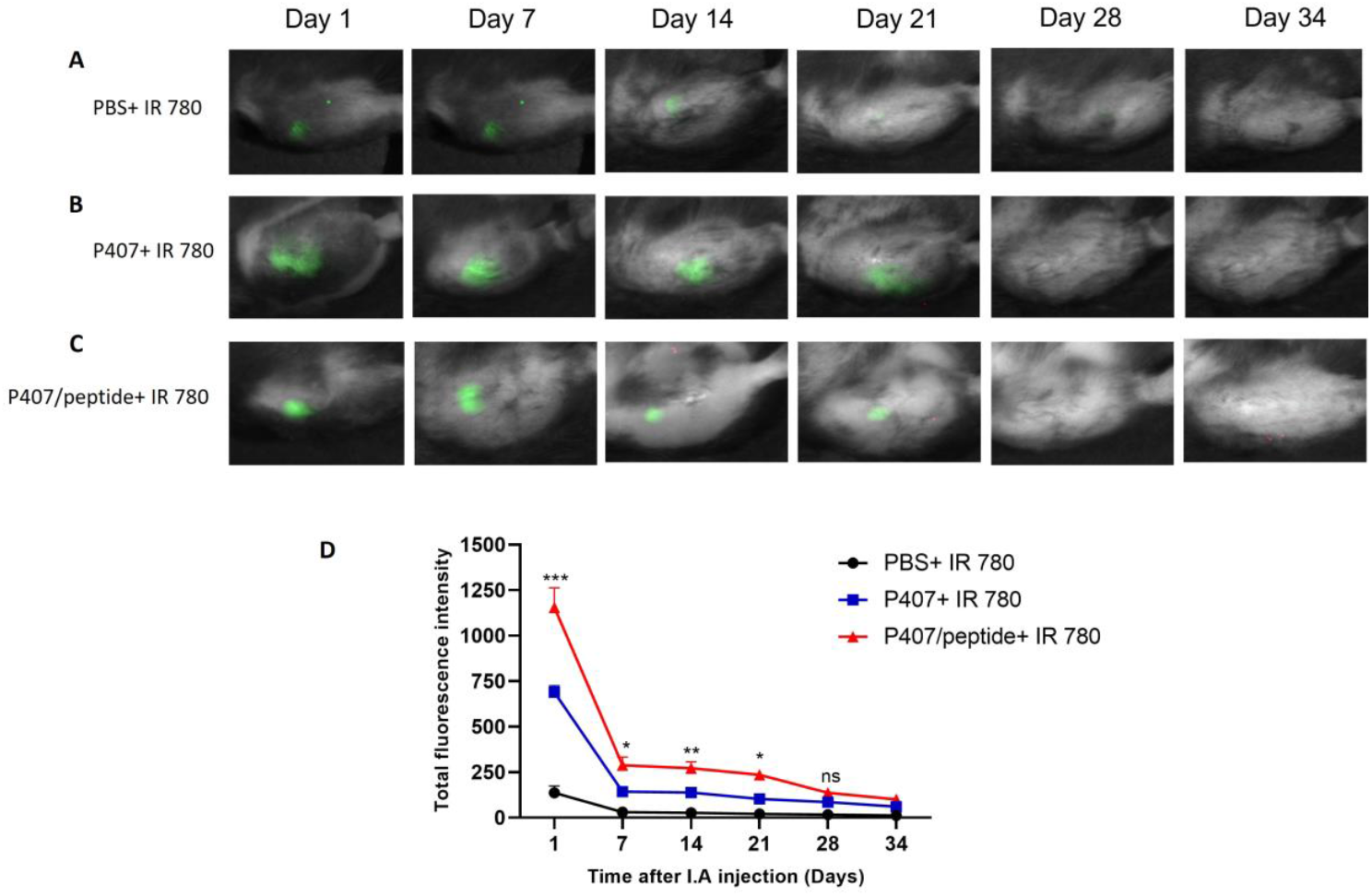
*In vivo* near-infrared fluorescence (NIRF) imaging and quantitative analysis of i.a injected formulations in the DMM mouse model. Representative NIRF images of IR780 fluorescence in the right knee joints of mice treated with (**A**) PBS + IR780, (**B**) P407 + IR780, or (**C**) P407-self-assembling peptide + IR780, captured on days 1, 7, 14, 21, 28, and 34 post-injection. (**D**) Quantification of total fluorescence intensity over time reveals prolonged retention of IR780 in the joint when delivered via P407 + self-assembling peptide hydrogel, compared to PBS or P407 alone. Data are presented as mean ± SD (n = 6). Statistical significance was determined by appropriate tests and is indicated as ***p ≤ 0.001, **p ≤ 0.01, *p ≤ 0.05, and ns = not significant. i.a.= Intra.articular

Quantitative analysis of total fluorescence intensity is shown in **Figure 5D**. On day 1, both hydrogel groups demonstrated significantly higher fluorescence intensities compared to the PBS group (p ≤ 0.001). Between days 7 and 21, fluorescence signals from the hydrogel groups remained significantly elevated (p ≤ 0.05 or p ≤ 0.01), with the P407/peptide hydrogel group consistently showing slightly higher retention than the P407 group. By day 28, differences between groups were no longer statistically significant, as the fluorescence signals gradually declined.

Fluorescence signals in all groups, although greatly reduced, remained detectable up to day 34. These findings align with the *in vitro* release profiles, where IR780 loaded into the P407–self-assembling peptide hydrogel exhibited a slower and more sustained release compared to P407 hydrogel or free dye.

### 3.6. Cell viability

The cytocompatibility of the 25% (w/v) P407–self-assembling peptide hydrogel was assessed using human chondrocyte (C28/I2) cells through MTS assay and LIVE/DEAD staining. C28/I2 cells were incubated with different concentrations of the P407–peptide hydrogel, and cell viability was evaluated after 24, 48 and 72 hours using the MTS assay.

As shown in **Figures 6A–C**, no decrease in cell viability was observed across different hydrogel concentrations, and no significant differences were detected between cultured cells. These results indicate that the P407–self-assembling peptide hydrogel is non-toxic and well tolerated by human chondrocytes.

**Figure 6:**
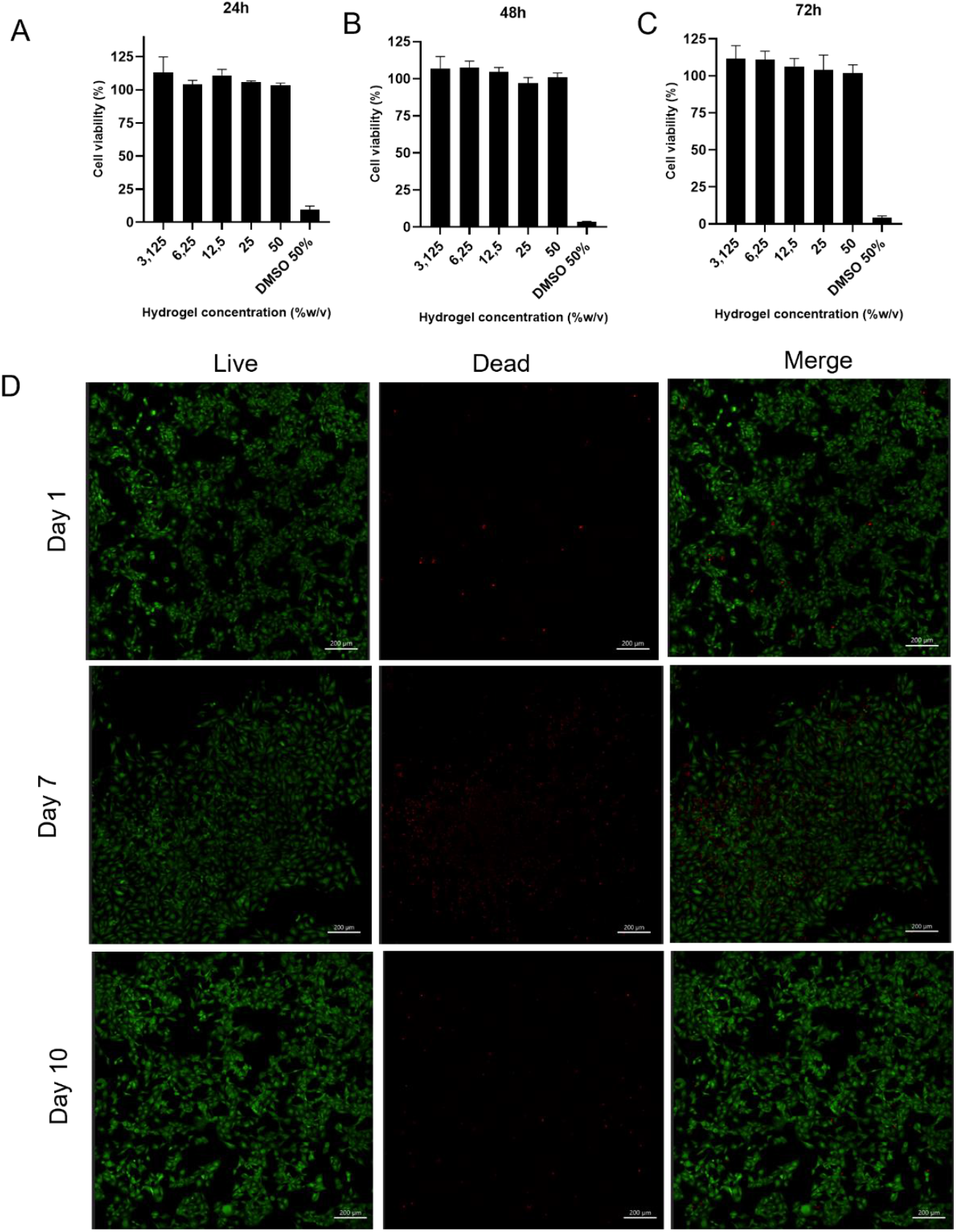
Evaluation of C28/l2 cell viability exposed to hydrogel using MTS assay. C28/I2 cells (A, B and C) were incubated for 24, 48 and 72 h with increasing concentrations of P407-self-assembling peptide up to a maximum of 50% (w/v). 50% DMSO in medium served as positive control. (D) LIVE/DEAD® staining of chondrocytes seeded on P407-self-assembling peptide hydrogels after 3, 7 and 10 days in culture. Scale bar = 200μm.

Further evaluation by LIVE/DEAD staining (**Figure 6D**) confirmed these findings. A high proportion of viable cells (green fluorescence) was observed at days 1, 3, and 7 of culture, with only few dead cells (red fluorescence) detected. The cells were homogeneously distributed across the hydrogel surface, demonstrating good cell attachment and viability over time. Moreover, an increase in the cell population was observed between days 7 and 10, indicating that the hydrogel not only supported cell survival but also promoted the adhesion and proliferation of C28/I2 cells.

These results demonstrate the excellent cytocompatibility of the 25% (w/v) P407–self-assembling peptide hydrogel, supporting its further evaluation for i.a. drug delivery applications.

## 4. Discussion

In this study, we developed a thermosensitive, injectable hydrogel composed of 25% (w/v) P407 and 1% (w/v) of a self-assembling peptide (Palmitoyl-WKGNNQQNYQQ), designed to enhance the physicochemical properties of the system for intra-articular applications.

P407 is a biocompatible triblock copolymer that exhibits thermosensitive behavior, facilitating rapid sol–gel transition at physiological temperatures and making it an attractive material for injectable drug delivery systems [23]. However, P407 hydrogels rapidly dissolve in aqueous environments, which limits their utility for sustained applications[24]. Although increasing the polymer concentration can improve gel firmness and delay dissolution, higher concentrations also result in elevated viscosity at room temperature, causing practical difficulties such as needle clogging during injection[25, 26]. To overcome these challenges, we incorporated a self-assembling peptide with sequence of Palmitoyl-WKGNNQQNYQQ into the P407 hydrogel formulation. The peptide’s amphiphilic design, incorporating hydrophobic residues (palmitoyl, tryptophan, tyrosine) and hydrophilic amino acids, was intended to reinforce the hydrogel structure through micellar interactions, thereby enhancing mechanical stability without compromising injectability.

We first evaluated the gelation behavior of the formulations at various concentrations of P407 with 1% of self-assembling peptide to assess their injectability and solidification properties. An ideal injectable hydrogel should remain fluid at room temperature for ease of handling and rapidly gel at body temperature to prevent undesired drug dispersion following injection[27]. Based on preliminary gelation tests, a 25% (w/v) P407 concentration was selected as it provided a balance between adequate gel strength and acceptable injectability, without causing needle clogging during administration. Higher P407 concentrations (>30%) resulted in excessively viscous solutions that were difficult to inject, while lower concentrations (<20%) failed to achieve reliable gelation at physiological temperature. Similarly, 1% (w/v) peptide was identified as the optimal additive concentration, as higher peptide loadings (>1%) led to the formation of bulk solid aggregates that impaired injectability Our results demonstrated that incorporation of 1% peptide into the 25% P407 formulation accelerated gelation at physiological temperature, reducing the sol–gel transition time from 44 to 32 seconds while maintaining good fluidity at room temperature. Such behavior improves handling characteristics, prevents premature needle clogging, and supports rapid in situ gel formation, facilitating localized drug depot creation upon administration[28]. Mechanical stability, evaluated by rheological analysis, further confirmed the reinforcing effect of the peptide. The peptide-modified hydrogel displayed a higher storage modulus and a slightly reduced gelation temperature compared to 25%P407 alone, indicating improved thermosensitivity and enhanced mechanical strength.

Moreover, the 25% modified hydrogel with peptide exhibited enhanced structural stability and a slower degradation rate relative to the same concentration of the P407 hydrogel alone. *In vitro* degradation studies revealed complete dissolution of the unmodified P407 hydrogel within 7 days, whereas the peptide-enhanced formulation remained intact for 9 days. This prolonged stability is particularly beneficial for intra-articular drug delivery, where extended hydrogel presence supports sustained release and tissue interaction. The slower degradation observed in the peptide-containing hydrogel is likely attributable to stabilizing molecular interactions — hydrogen bonding, hydrophobic association — between the peptide nanofibers and P407 chains, which reduce water penetration and delay matrix erosion. Mechanistic insights support these observations: in deionized water, the peptides formed stable β-sheet structures, while exposure to electrolyte solutions triggered self-assembly into interwoven nanofibrous networks[29, 30]. These nanostructures enhanced both the mechanical integrity and the resistance to dissolution of the hydrogel, aligning with the observed prolongation of gel presence.

The improved degradation and stability profiles were further validated *in vivo* using a DMM mouse model of OA. *In vivo* results demonstrated that the degradation of the P407-peptide hydrogel was partial and occurred slowly, consistent with findings from the *in vitro* degradation study. The preferability of slow degradation in injectable hydrogels lies in the ample time it affords for the integration of loaded drugs or cells into nearby tissue and the facilitation of natural cartilage repair as the hydrogel gradually breaks down[31].

In summary, this novel. thermosensitive. injectable hydrogel. engineered through the incorporation of the self-assembling peptide into the poloxamer matrix. represents a significant advancement in the field of regenerative medicine. Its rapid gelation kinetics, biocompatibility, and capacity for localized drug delivery make it an attractive platform for addressing a wide spectrum of tissue regeneration challenges, with particular relevance to musculoskeletal disorders such as OA. Our research sets the stage for potential breakthroughs in therapeutic strategies aimed at enhancing tissue repair and regeneration.

## Acknowledgments

The research leading to these results has received funding from the European Union’s Horizon 2020 research and innovation program AutoCRAT under grant agreement No 874671. The material presented and views expressed here are the responsibility of the author(s) only. The EU Commission takes no responsibility for any use made of the information set out.

## Author Contributions

S.S.S and L.J.C designed the experiments. S.S.S., S.M.L., and S.R. performed the *in vitro* experiments. S.S.S. and T.S. performed the *in vivo* experiments. S.S.S. was responsible for most of the data and statistical analysis and drafting of the manuscript. All authors have read, revised and agreed to the published version of the manuscript.

## Declaration of Transparency and Scientific Rigour

This Declaration acknowledges that this paper adheres to the principles for transparent reporting and scientific rigor of preclinical research as stated in the BJP guidelines for Design & Analysis, and Animal Experimentation, and as recommended by funding agencies, publishers, and other organizations engaged with supporting research.

## Informed Consent Statement

Not applicable.

## Conflict of Interest

The authors declare no potential conflicts of interest.

## Data Availability Statement

The original contributions presented in the study are included in the article/supplementary material. Further inquiries can be directed to the corresponding author.

## Conflicts of Interest

No conflict of interest exists in the submission of this manuscript, and the manuscript is approved by all authors for publication

## Supplementary material

**Supplementary Figure 1:**
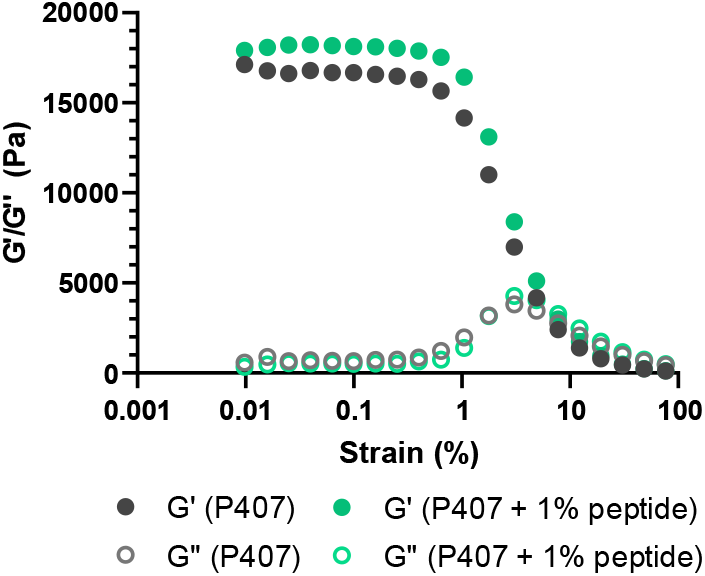
Strain sweep of 25% P407 and 25% P407 -peptide hydrogels. G’ and G’’ were measured as strain was increased from 0.01% to 100% at a constant frequency of 1 Hz. A linear viscoelastic regime was observed for the strain range between 0.01% and 0.4%.

**Supplementary Figure 2:**
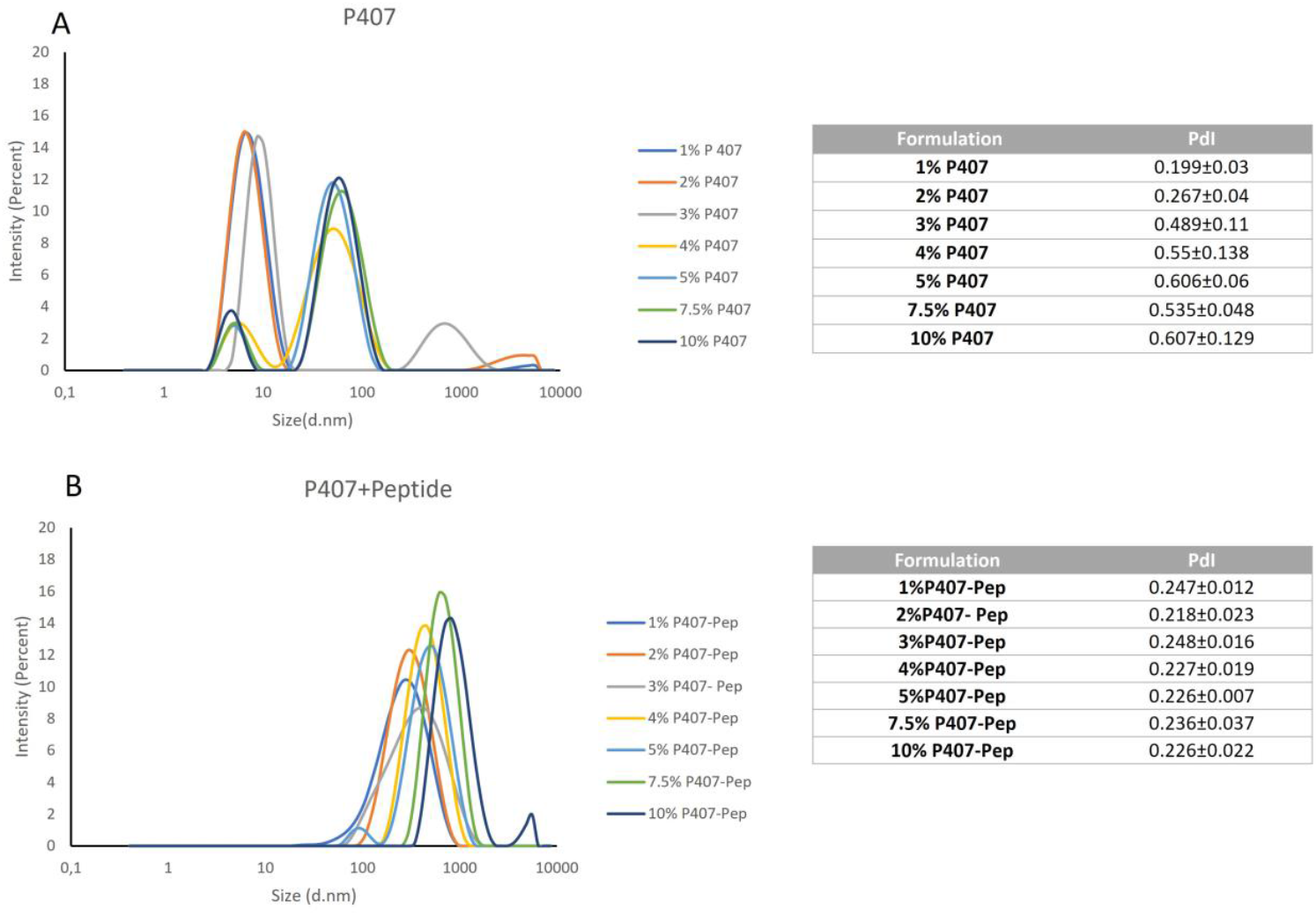
DLS measurements with representative histogram with the pick size distribution and PdI valus for (**A**) 1, 2, 3, 4, 5, 7.5 and 10% (w/w) P407. (**B**) P407-peptide. DLS= Dynamic Light Scattering; PdI= Polydispersity Index.

## Notes

### Competing Interest Statement

The authors have declared no competing interest.

